# Comparative analysis of nuclei isolation methods for brain single-nucleus RNA sequencing

**DOI:** 10.1101/2025.03.25.645306

**Authors:** Holly N. Kersey, Dominic J. Acri, Luke C. Dabin, Kelly Hartigan, Richard Mustaklem, Jung Hyun Park, Jungsu Kim

## Abstract

Single-nucleus RNA sequencing (snRNA-seq) enables resolving cellular heterogeneity in complex tissues. snRNA-seq overcomes limitations of traditional single-cell RNA-seq by using nuclei instead of cells, allowing to utilize frozen tissues and difficult-to-isolate cell types. Although various nuclei isolation methods have been developed, systematic evaluations of their effects on nuclear integrity and subsequent data quality remain lacking, a critical gap with profound implications for the rigor and reproducibility. To address this, we compared three mechanistically distinct nuclei isolation strategies with brain tissues: a sucrose gradient centrifugation-based method, a spin column-based method, and a machine-assisted platform. All methods successfully captured diverse cell types but revealed considerable protocol-dependent differences in cell type proportions, transcriptional homogeneity, and the preservation of cell-type-specific and cell-state-specific markers. Moreover, isolation workflows differentially influenced contamination levels from ambient, mitochondrial, and ribosomal RNAs. Our findings establish nuclei isolation methodology as a critical experimental variable shaping snRNA-seq data quality and biological interpretation.

**MOTIVATION:** Single-nucleus RNA sequencing (snRNA-seq) has become an essential tool for transcriptomic analysis of complex tissues. However, the quality and efficiency of data generation depend heavily on the method used for nuclear isolation. The existing isolation techniques vary in their ability to preserve nuclear integrity, minimize ambient RNA contamination, and optimize recovery rates. Despite these differences in quality, a systematic comparison of these methods, specifically for brain tissue, is lacking. This gap poses a challenge for researchers in choosing the most suitable approach for their particular experimental requirements. To address this critical issue, our study directly compared three nuclei isolation methods and evaluated their performance in terms of yield, purity, and downstream sequencing quality. By providing a comprehensive assessment, we aim to guide researchers in selecting the most appropriate isolation protocol for their snRNA-seq experiments, ensuring optimal results and advancing the study of complex brain tissues at the single-nucleus level.

## INTRODUCTION

Single-nucleus RNA sequencing (snRNA-seq) is an effective approach for investigating gene expression at the single-cell level, particularly in complex tissues such as the brain[1-4]. This technique addresses the key limitations of traditional single-cell RNA sequencing (scRNA-seq), which can become problematic when applied to brain tissue[5-7]. For example, conventional cell dissociation protocols often compromise the integrity of neurons through mechanical shearing or induce the activation of glia[6, 8-12]. These factors contribute to cell-type isolation bias, the potential loss of rare or fragile cell populations, and *ex vivo* gene expression artifacts[13]. snRNA-seq overcomes the need for intact cell dissociation, thereby minimizing dissociation-associated challenges and providing a more physiologically accurate representation of the brain’s cellular heterogeneity[12, 14, 15]. Additionally, the use of nuclei, rather than whole cells, enables the study of samples that were previously difficult to analyze via scRNA-seq, such as postmortem, frozen, or biobank tissue, including those obtained from individuals with neurodegenerative conditions such as Alzheimer’s disease (AD)[12, 16]. This advancement opens new avenues for investigating cellular and molecular changes in complex disorders, potentially leading to improved understanding of disease mechanisms and the development of targeted therapies[17].

Given the benefits of snRNA-seq, assessing and improving the quality of preclinical snRNA-seq datasets has become a growing research priority. Recent efforts in single-cell biology have yielded comprehensive atlas-level datasets, such as the Human Cell Atlas, BRAIN Initiative Cell Consensus Network, and Allen Brain Atlas[18-21]. These and other single-cell initiatives have uncovered potential disease mechanisms and identified new drug targets [22-26]. Funding agencies, such as the National Institutes of Health (NIH) and Chan Zuckerburg Initiative (CZI), have invested heavily in single-cell research to advance our understanding of human health, disease, and treatment[27-31]. snRNA-seq can further enhance these efforts, offering unique advantages in terms of tissue compatibility.

However, the potential of snRNA-seq in advancing single-cell genomics critically depends on the quality of nuclear preparations. Therefore, there is an urgent need to preserve high-quality nuclei preparations to ensure that valuable samples and substantial sequencing costs are not wasted due to technical artifacts or contamination by ambient RNAs that necessitate resequencing. Our study addresses this critical need by comparing nuclei isolation protocols to identify methods that best maintain nuclear integrity and purity throughout the snRNA-seq workflow.

Low-quality nuclei isolation protocols can introduce artifacts that compromise the proper data interpretation and the experimental reliability[32, 33]. Contaminants, such as ambient RNA, ribosomal RNA, and mitochondrial RNA, can cause technical artifacts that mask biological effects. For example, ambient RNA from lysed cells, a ubiquitous issue of scRNA- and snRNA-seq datasets, can lead to false positives in gene expression profiles, obscuring true cell type-specific markers[34, 35]. Excessive levels of contaminating mitochondrial and ribosomal RNA can overwhelm sequencing data, reducing the depth of informative transcripts and hindering the detection of low-abundance genes[36]. Furthermore, such artifacts can significantly compromise various aspects of downstream analyses, including accurate cell type identification, differential expression analysis, and integration with other omics information. While computational tools can mitigate some variability in sample preparation, obtaining high-quality nuclei isolations before library preparation is essential for minimizing technical noise and ensuring reliable, reproducible data[37-40].

Isolating nuclei from complex tissues such as the brain poses challenges in preserving cellular diversity, minimizing RNA degradation, and reducing extranuclear contamination. To address these challenges, we compared three mechanistically distinct nuclei isolation protocols: (1) manual homogenization followed by sucrose gradient centrifugation, (2) a spin column-based method, and (3) a machine-assisted platform[41-43]. Each nuclei isolation method for brain tissue processing has distinct advantages and limitations. The sucrose gradient centrifugation method is a well-established and cost-effective technique. However, this method can suffer from person‒to-person variability in hand grinding and gradient preparation and may require ultracentrifugation. The column-based method provides comparable scalability without the need for specialized machinery, providing faster processing times than sucrose gradient preparation. However, this protocol still faces potential variability in tissue grinding and requires specific consumable columns. The machine-assisted platform provides an automated approach that reduces processing time and minimizes sample-to-sample and person-to-person variability. However, it requires the purchase of specialized equipment and specific consumable cartridges. Notably, all three methods typically involve enzymatic digestion, which can potentially activate certain cell types, introducing a common source of potential bias[10]. The choice of method ultimately depends on factors such as available equipment, sample size, desired throughput, and tolerance for variability, requiring researchers to carefully consider these aspects when selecting a nuclei isolation approach for their specific experimental needs.

Using the mouse brain cortex, we assessed the efficacy of these methods by preparing nuclei suspensions, capturing RNA using 10x Genomics’ Chromium, and sequencing snRNA-seq libraries. Given the heterogeneity of brain tissue, we sought to preserve various neuronal and glial populations while minimizing bias and artifacts. Each isolation method was evaluated for nuclei yield, contamination, and marker gene expression profiles. Our comparative analysis provides valuable insights into nuclei isolation strategies for brain tissues, enhancing our ability to study complex neurological processes and disorders at the single-nucleus level.

## RESULTS

### Nuclei yield and viability differ across isolation protocols

We evaluated three distinct nuclei isolation protocols optimized for single-nucleus RNA sequencing: manual homogenization followed by sucrose gradient centrifugation, a commercially available spin column-based method, and a machine-assisted platform (**Figure 1A**). All isolations were performed using cortical tissue from 6-month-old C57BL/6J mice (N=2/method). Cortical tissue was dissected from the anterior portion of one hemisphere, and approximately 30 mg was weighed to normalize the input for each method. Representative brightfield microscopy images revealed visible differences in preparation quality among the methods. The centrifugation-based method produced defined individual nuclei with minimal background debris (**Figure 1B)**. In contrast, the column-based method resulted in densely packed nuclei with notable aggregation and substantial debris contamination even after multiple optimization attempts (**Figure 1C**). The machine-assisted method yielded well-separated, intact nuclei with negligible debris (**Figure 1D**). Among the 6 samples selected for sequencing, the centrifugation-based and machine-assisted methods provided similar yields of 2 million nuclei or approximately 60,000 nuclei per milligram of input (**Figure 1E**). In contrast, the column-based approach yielded 25% fewer nuclei with similar input material. Of the total number of nuclei recovered, the machine-assisted method maintained the structural integrity of almost 100% of the nuclei. This result indicates that the extracted nuclei remained intact upon recovery (**Figure 1F**). The centrifugation-based method preserved 85% of nuclei in their intact form, whereas the column-based method yielded only 35% structurally intact nuclei. These findings demonstrate that, compared with the column-based approach, the centrifugation-based and machine-assisted methods are more effective in terms of both nuclear yield and purity.

**Figure 1:**
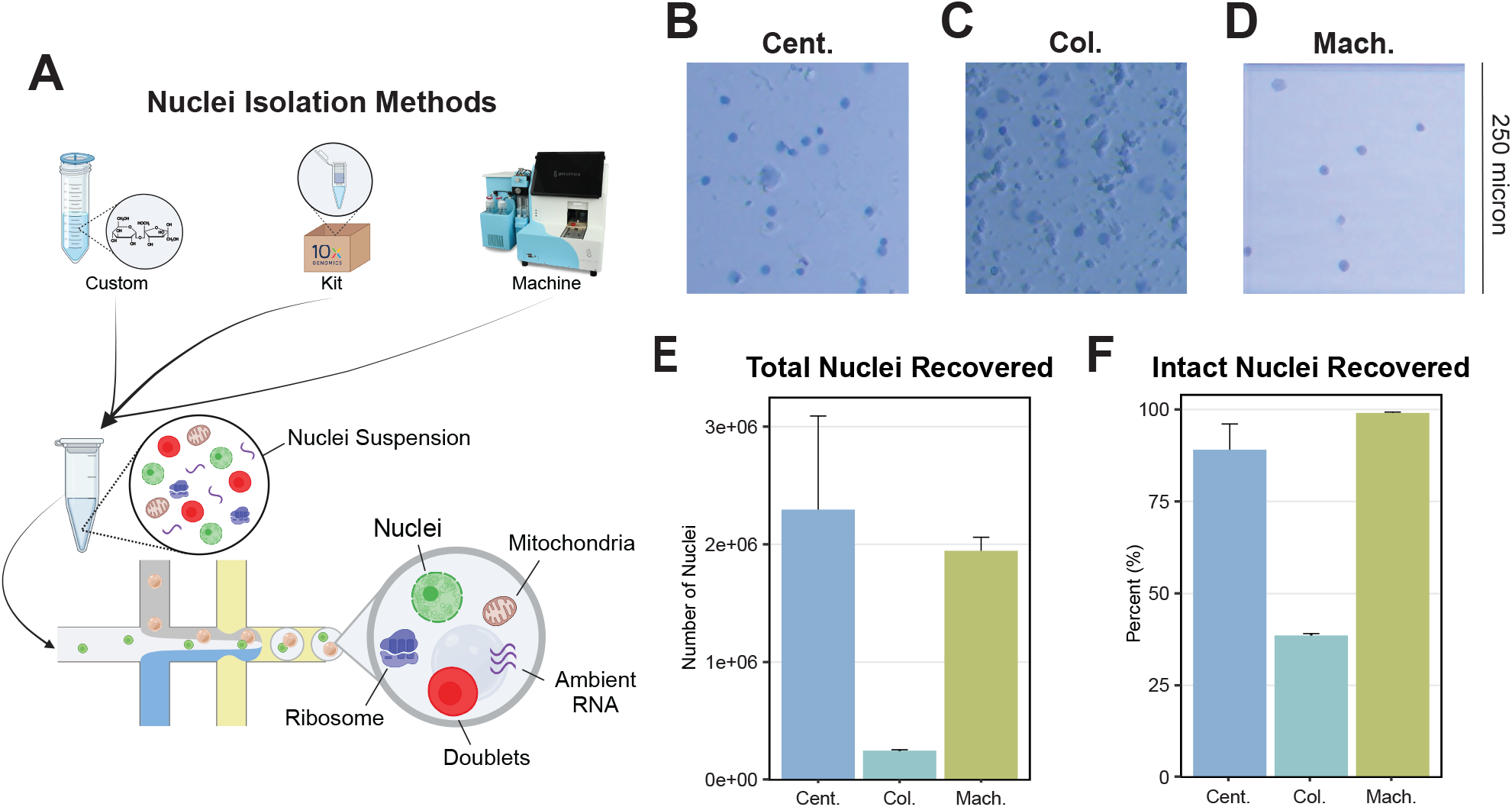
Nuclei yield and viability differ across isolation protocols. **(A)** Graphical abstract depicting the workflow of each method and highlighting potential contaminants, including mitochondria, ribosomes, and ambient RNA, which can be captured in droplet-based sequencing depending on the isolation approach. Representative images of trypan blue-stained nuclei demonstrating the morphology and quality of nuclei obtained from the **(B)** centrifugation-based (Cent.), **(C)** column-based (Col.) and **(D)** machine-assisted (Mach.) isolation methods. The scale bar represents 250 microns. (**E**) The total number of nuclei recovered from each method, as quantified using the Countess Automated Cell Counter, reveals differences in yield among the protocols. (**F**) From the total nuclei population isolation, the proportion of structurally intact nuclei recovered from each method. Error bars represent standard deviation.

### Isolation technique influences the cell types captured in snRNA-seq

Next, we performed snRNA-seq experiments to evaluate the impact of the different isolation methods on the transcriptional profiles using 10x Genomics’ Chromium single-cell 3’ gene expression assay. In total, 98,452 nuclei were captured from the six samples, forming 36 distinct clusters (**Figure 2A**). The defined clusters included astrocytes (A), microglia (M), excitatory neurons (eN), inhibitory neurons (iN), oligodendrocytes (O), and other cell types (Ot). Clusters were annotated using the single-cell Mouse Cell Atlas (scMCA) (**Supplementary Table 1**) and further refined with PanglaoDB [44, 45]. Key marker genes exhibited distinct expression patterns across cell types (**Figure 2B**). Notably, *Gja1* and *Slc1a3* were strongly expressed in astrocyte clusters, whereas *Sv2b* and *Slc17a7* were robustly expressed across excitatory neuron populations. *Gad1* and *Gad2* expressions were restricted to inhibitory neuron clusters. Across all methods, most of the nuclei captured were excitatory neurons (53.9%), followed by inhibitory neurons (17.2%) (**Figure 2C**). Glial cells and other cell types comprised smaller proportions of the total population, with mean percentages varying according to the isolation method. Interestingly, the centrifugation-based method captured the largest proportion of astrocytes (13.9%), whereas the machine-assisted method attained the largest proportions of microglia (5.6%) and oligodendrocytes (15.9%). The uniformity of identified cell populations is a crucial quality control matix for accurate cell type identification. To evaluate this, population homogeneity was assessed via the ratio of global unshifted entropy (ROGUE), a metric that quantifies the transcriptional consistency within cell populations (**Figure 2D**)[46]. Most cell types exhibited moderate homogeneity, with ROGUE values between 0.5 and 0.75. In particular, astrocytes displayed elevated ROGUE values (>0.72) across all three methodologies, indicating highly uniform transcriptional profiles within this cell population. In all other cell types, both the machine-assisted and column-based methods produced comparable scores, between 0.65 and 0.8, whereas the centrifugation-based method yielded lower ROGUE values, <0.68, implying less uniformity within these populations. These data demonstrate that although all three nuclear isolation methods successfully capture diverse cell types from mouse cortical tissue, they exhibit differences in cell type proportions and transcriptional homogeneity.

**Figure 2:**
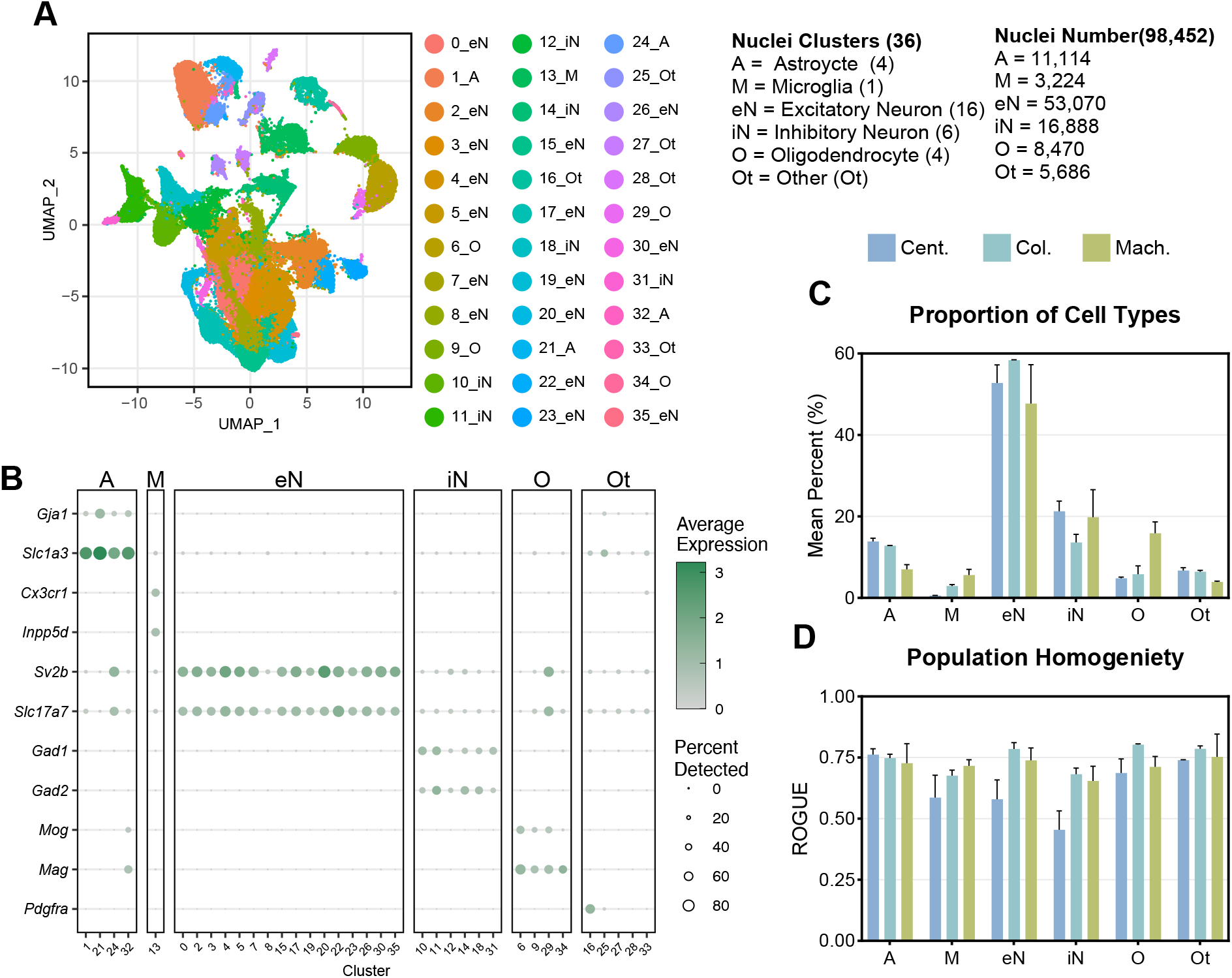
Isolation technique influences the cell types captured in snRNA-seq. **(A)** Uniform Manifold Approximation and Projection (UMAP) plot of 98,452 nuclei including astrocyte (A), microglia (M), excitatory neurons (eN), inhibitory neurons (iN), oligodendrocytes (O), and other (Ot) clusters. **(B)** Dot plot of key marker genes for each cell type. Average scaled expression level scales with white (low) to green (high) shading. The size of dots represents the percent of nuclei in each cell type expressing the gene. **(C)** The mean percent of each type of nuclei captured relative to the total population for each nuclei isolation method. **(D)** The homogeneity of nuclei populations as reported by the ROGUE value, an entropy-based statistic that measures the uniformity of given populations, using ROGUE. Color represents the isolation method (centrifugation-based=Cent., column-based=Col., machine-assisted=Mach.). Error bars represent standard deviation.

### Differences in Quality Control Metrics Among Isolation Protocols

We evaluated the quality of the nuclear preparations using standard metrics commonly employed in sn- and scRNA-seq experiments. We predicted the presence of doublets, which are technical artifacts where mRNAs from two different nuclei are tagged with the same barcode and cannot be demultiplexed, using DoubletFinder. DoubletFinder uses a subset of a scRNA-seq dataset to generate artificial doublets, combines them with the remaining data, and employs principal component analysis (PCA) to measure the proportion of artificial nearest neighbors for each cell, thresholding these values to produce final doublet predictions[38]. The column-based method yielded inconsistent doublet rates among samples, with predicted percentages ranging from a low of 3.2% to a high of 5.7% (**Figure 3A**). In the context of snRNA-seq analysis, a consistent doublet rate is generally preferable as it indicates more reliable and reproducible sample processing. The remaining samples produced by the centrifugation-based and machine-assisted methods had comparable doublet rates, ranging between 4.5% and 5.5%. To quantify ambient RNA contamination, we utilized SoupX, which calculates the contamination fraction parameter “rho”. This parameter estimates the proportion of unique molecular identifiers (UMIs) attributable to ambient RNA by analyzing empty droplets and determining cell-specific contamination levels[37]. This approach identifies and removes nuclei-free RNAs, which can confound the biological interpretation of single-nuclei transcriptomic data. Column-based samples presented significantly elevated rho values (>0.25), nearly twofold higher than those of both centrifugation-based and machine-assisted methods, indicating substantial ambient RNA contamination (**Figure 3B**). The machine-assisted method again displayed consistency, with rho values of approximately 0.14 for both samples. The samples produced via the centrifugation-based method presented some degree of intersample variability, with rho values between 0.07 and 0.17. Recent studies have suggested that *Malat1* (metastasis-associated lung adenocarcinoma transcript 1), a long noncoding nuclear RNA, can act as a quality metric for snRNA-seq, with low levels identifying nuclei of poor quality[47, 48]. There was no discernable difference between *Malat1* expression, with all samples having median expression levels, the log-transformed RNA counts, of approximately 5 (**Figure 3C**).

**Figure 3:**
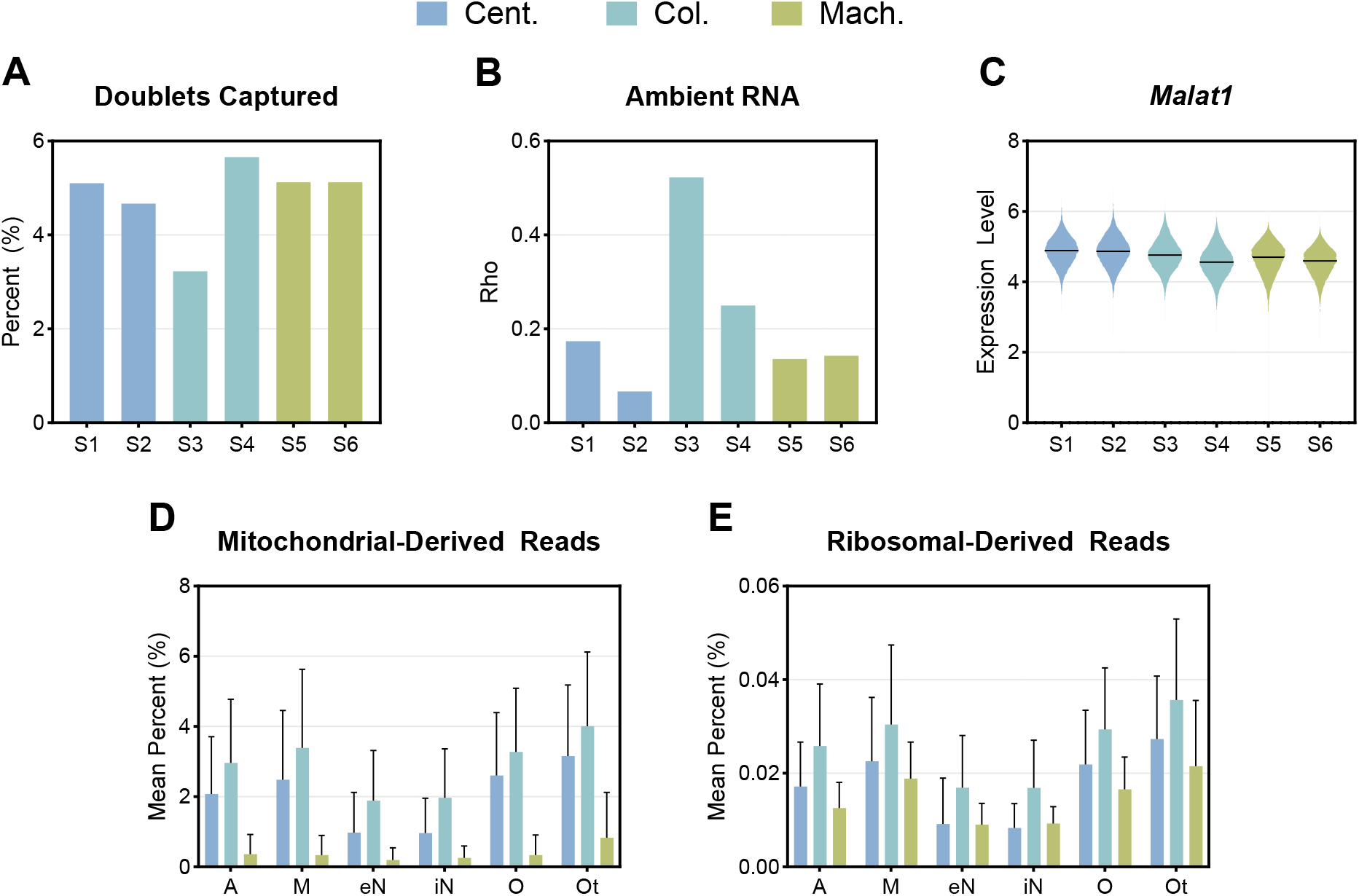
Differences in Quality Control Metrics Among Isolation Protocols. **(A)** The percentage of nuclei identified as doublets for each sample using DoubletFinder, representing the proportion of doublets relative to the total number of captured nuclei. **(B)** The contamination fraction as reported by Rho value, an indicator of ambient RNA contamination, for each sample using SoupX. **(C)** Violin plot of *Malat1*, a marker of nuclear integrity, expression for each sample. The line represents the median expression. The mean percentage of **(D)** mitochondrial-derived reads and **(E)** ribosomal-derived reads in each cell type per isolation method. Error bars represent standard deviation. Color represents the isolation method (centrifugation-based=Cent., column-based=Col., machine-assisted=Mach.).

High-quality nuclei preparations should contain minimal to no cytoplasmic content. In snRNA-seq, the presence of non-nuclear material can introduce bias in gene expression profiles and compromise the accuracy of downstream analyses. Therefore, we analyzed mitochondrial and ribosomal transcript levels as indicators of cytoplasmic contamination. Importantly, the machine-assisted method demonstrated low levels of average mitochondrial-derived reads, <0.5% of total reads, across all cell types (**Figure 3D**). The centrifugation- and column-based methods resulted in elevated average levels of mitochondrial-derived reads, ranging from 1–3% and 2–4%, respectively. Although ribosomal-derived reads remained minimal across all preparation methods, the machine-assisted and column-based protocols displayed the lowest contamination, with values < 0.03% (**Figure 3E**). The column-based method demonstrated the highest levels of contamination, with ribosomal-derived reads exhibiting an average contamination level of 0.036%. The results indicate that the machine-assisted method yielded nuclei preparations with high consistency between samples and low contamination levels, whereas the column-based and centrifugation-based methods presented greater sample-to-sample variability and elevated contamination levels.

### Method-specific variations in glial gene expression

The accurate identification and characterization of specific cell types, particularly glial cells, is crucial for understanding brain function and pathology in single-cell genomic studies. Different nuclei isolation methods may introduce biases or artifacts that affect the detection of cell type-specific markers, potentially leading to misclassification or incomplete representation of cellular populations. Therefore, we analyzed the gene expression profiles across the three isolation methods, with a focus on glial cell types. We compared markers for the microglia population identified by each isolation method with various glial cell type annotations from the Panglao database (**Supplementary Table 2**)[45]. To address the specificity of these markers, we assessed the proportion of identified microglia markers by each method that corresponds to different glial cell types. Notably, both the centrifugation-based and machine-assisted methods resulted in significant enrichment (p value < 0.05) of microglial markers in the Panglao database, with the machine-assisted method yielding the highest proportion of microglia-specific markers (**Figure 4A**). Further investigation of canonical microglia-specific markers revealed method-specific variations in expression patterns. For example, *C1qa* exhibited the highest expression with the machine-assisted method (**Figure 4B**). Two other microglia-specific markers, *Hexb* and *Siglech* showed robust expression across all methods but was greater expression in the centrifugation-based and machine-assisted methods (**Figure 4C, D**). Interestingly, *Tmem119* expression was elevated specifically in the machine-assisted method (**Figure 4E**). Despite the enrichment of microglia-specific markers in microglia nuclei, some of them may express markers for other glial cell types, such as oligodendrocytes and satellite glial cells. When examining oligodendrocyte markers within the microglial population, the machine-assisted method showed negligible expression of *Plp1*, whereas both the centrifugation- and column-based methods exhibited some level of *Plp1* expression (**Figure 4G**). Only the centrifugation-based method demonstrated the expression of *Mbp* (**Figure 4H**). Similarly, analysis of satellite glial markers within microglia nuclei revealed the highest expression of *Glul* and *Ptgds* with the centrifugation-based method (**Figure 4, I, J**). Taken together, our findings highlight critical method-specific strengths and limitations. For example, machine-assisted isolation maximizes microglia-specific signals while minimizing oligodendrocyte marker contamination. The observed technical variability underscores the importance of isolation method selection in glial cell research. Method-driven differences in marker detection could significantly impact interpretations of cellular identity and function, particularly in studies comparing glial subpopulations or analyzing rare cell types.

**Figure 4:**
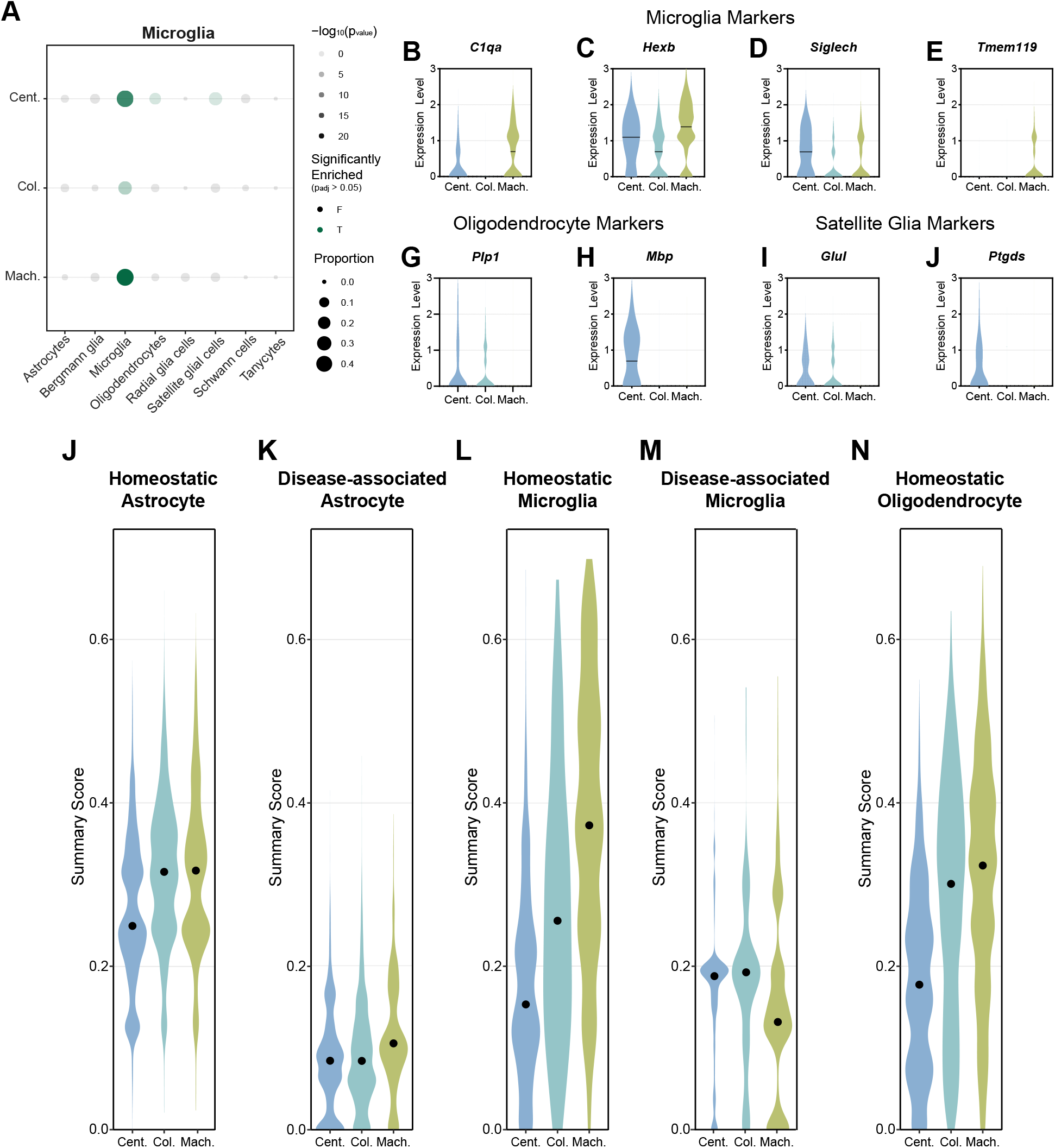
Method-specific variations in glial gene expression. **(A)** Dotplot of identified microglia marker gene expression compared to glial cell type annotations in the Panglao database. The size of the dots represents the proportion of microglia marker genes expressed within each glial cell type. The shading of dots represents the -log_10_(p_value_) of the marker gene expression in the glial cell type and the color of dots represents if the expression is significant (green) or not significant (black) determined by Fisher’s exact test. Violin plots of expression level of canonical microglia markers, **(B)** *C1qa*, **(C)** *Hexb*, **(D)** *Siglech*, and **(E)** *Tmem119*, as well as oligodendrocyte markers, **(F)** *Plp1* and **(G)** *Mbp*, and satellite glia markers, **(H)** *Glul* and **(I)** *Ptgds*, within the identified microglia population. The line represents the median expression level. UCell summary scoring of cell states, including **(J)** homeostatic astrocyte, **(K)** disease-associated astrocyte, **(L)** homeostatic microglia, **(M)** disease-associated microglia, and **(N)** homeostatic oligodendrocyte, within their respective cell type population. Dots represent the median summary score. Color represents the isolation method (centrifugation-based=Cent., column-based=Col., machine-assisted=Mach.).

Several key glial genes in our analysis have been implicated in various neurodegenerative conditions. For instance, *Tmem119* expression is significantly reduced in activated microglia in AD, whereas *C1qa* is associated with microglial activation after traumatic brain injury[49-51]. Also, a decrease in astrocytic *Glul* expression has been associated with epilepsy and subsequent neurodegeneration[52, 53]. Therefore, we explored gene signatures across various cell states, with UCell summary scoring strategy. This approach uses rank-based enrichment to calculate the relative expression of predefined gene sets within individual cells or nuclei (**Supplementary Table 3**)[54]. Compared with the centrifugation-based method, the column-based and machine-assisted methods yielded similar higher signature scores (approximately 0.3) for homeostatic markers in astrocytes (**Figure 4J**). The disease-associated astrocyte signature generally had low scores, with median scores <0.2. (**Figure 4K**). In microglia, the machine-assisted method had the highest scores for homeostatic signatures and the lowest scores for disease-associated signatures (**Figure 4L, M**). When homeostatic oligodendrocytes were examined, the column-based and machine-assisted methods demonstrated the highest signature scores, >0.3, with a notable reduction in the centrifugation-based method score, <0.2 (**Figure 4N**). Taken together, our comprehensive analyses demonstrate that the choice of nuclear isolation methods significantly affect gene expression patterns and cellular state signatures. Each method varies in its ability to preserve cell type-specific markers and homeostatic states across astrocytes, microglia, and oligodendrocytes, highlighting the importance of method selection in glial cell research.

## DISCUSSION

Selecting an appropriate nuclei isolation method is crucial for ensuring reliable and reproducible snRNA-seq data, especially for brain tissue[6]. The heterogeneous brain composition, comprising various neuronal subtypes, glia, and other specialized cells, presents unique challenges for accurate transcriptomic profiling. Moreover, the risk of *ex vivo* activation during sample preparation is a significant concern, as the gene expression of microglia and other cell types can be rapidly altered in response to environmental factors[10]. While researchers often select isolation protocols based on ease of use, sample availability, or throughput, there have been few systematic evaluations of how these choices impact data quality, cell type representation, and gene expression. Differences in nuclei isolation methods can introduce technical artifacts, leading to inconsistencies in data interpretation and limiting cross-study comparisons. Additionally, technical variability between samples can introduce confounding factors, making it difficult to discern true biological differences from methodological artifacts[55]. These issues also have implications for statistical power because increased technical variability may necessitate larger sample sizes to detect meaningful biological differences. To address these critical issues, we performed a comparative analysis of different isolation protocols and found significant differences in yield, quality, and transcriptional fidelity, all of which have direct implications for downstream analyses. With the growing trend towards expansive, collaborative, and data-driven research in the field, it becomes crucial to comprehend the methodological variations among datasets. This understanding is vital for maintaining reproducibility and ensuring biological accuracy, especially as researchers increasingly rely on large-scale analyses, integrative studies, and published datasets [56].

Our study highlights substantial variability in transcriptional profiles across different nuclei isolation methods. The centrifugation-based method yielded the greatest number of nuclei but also exhibited pronounced transcriptional heterogeneity, likely due to differences in RNA degradation or cytoplasmic contamination between individual preparations (**Figures 1, 2**). This suggests that, while maximizing yield, the centrifugation-based method may compromise data integrity by introducing technical noise in a sample-by-sample manner. In contrast, the machine-assisted method resulted in the highest nuclei viability and the lowest levels of mitochondrial and ribosomal RNA contamination (**Figures 1, 3**). These advantages are likely due to its automated processing, which reduces the handling time and minimizes the number of technical artifacts. Additionally, the machine-assisted method demonstrated greater consistency across samples in key quality control metrics, including doublet formation and ambient RNA contamination, whereas the centrifugation- and column-based methods exhibited high variability. Such variability can introduce batch effects, leading to inconsistencies in downstream analyses and unreliable biological conclusions[55, 57]. The column-based method, while yielding the lowest number of nuclei, also resulted in the highest levels of ambient RNA contamination. This contamination may distort transcriptomic profiles, leading to the underrepresentation of specific cell types or misclassification of cellular states[34, 37]. Given the importance of accurately identifying distinct cellular populations in neurodegenerative diseases such as AD, these limitations are particularly concerning. For example, misidentifying disease-associated microglia (DAMs), reactive astrocytes, or rare cell subpopulations can hinder efforts to characterize disease progression and cellular dysfunction in AD[58]. This, in turn, could lead to wasted effort and funding spent pursuing hypotheses based upon unreliable data.

Many cell types, such as microglia, are especially sensitive to dissociation-induced activation, which is a known challenge in single-cell studies[9, 10]. When microglial populations were examined, the machine-assisted method preserved the strongest expression of homeostatic microglial markers and the most accurate representation of their transcriptional signatures (**Figure 4**). Because the mice used in this study lacked disease pathology, we expected minimal expression of DAM-related genes. However, both the centrifugation- and column-based methods resulted in elevated DAM signatures, suggesting that technical artifacts, rather than true biological signals, may drive these transcriptional changes. Such artifacts can lead to erroneous classification of microglial states, potentially confounding our understanding of neuroinflammatory processes in AD and other neurodegenerative diseases. Accurate classification of homeostatic versus reactive microglia is critical for studying neurodegeneration, as misinterpretation of transcriptional profiles may obscure disease-specific changes and misdirect therapeutic development[59]. Our profiling experiments emphasize the importance of selecting an isolation method that minimizes technical distortions to ensure biologically meaningful conclusions.

These findings highlight that nuclear isolation methods may introduce technical artifacts that complicate direct comparisons between studies using different approaches. Each method significantly affects data quality, cell type representation, and transcriptional integrity. The observed variability in centrifugation- and column-based methods highlights the potential risks of relying on high-yield or commercial protocols without fully considering their impact on downstream analyses. This variability not only compromises data quality, cell type representation, and transcriptional integrity but also underscores the need for methodological transparency and rigorous validation. By demonstrating how method selection can alter transcriptional signatures, our study emphasizes the importance of adhering to FAIR guidelines, i.e., making data Findable, Accessible, Interoperable, and Reusable (FAIR Principles)[60]. By ensuring that data are both interoperable and reusable, researchers can more easily replicate and validate each other’s findings, addressing the reproducibility crisis in scientific research. The ability to integrate and reuse data from diverse sources has accelerated the discovery of new biological insights, ultimately advancing our understanding of cellular diversity and disease mechanisms. The reliability of the machine-assisted method in maintaining low cytoplasmic contamination across samples provides a model for optimizing high-throughput and large-scale studies. In contrast, the inconsistency observed in other methods highlights the need for protocol refinement to ensure accurate biological representation. By demonstrating that method selection can alter transcriptional signatures, our study emphasizes the necessity of methodological transparency and careful validation when datasets from different studies and conditions are compared. Future research should focus on refining isolation strategies across different brain regions, species, and pathological conditions to ensure the most accurate representation of cellular diversity and disease mechanisms at the single-cell level. Optimizing nuclei isolation protocols can improve the accuracy and reproducibility of snRNA-seq studies, ultimately advancing our understanding of cell diversity and disease mechanisms at the single-cell level.

## LIMITATIONS OF THE STUDY

This study provides valuable insight into methods for isolating nuclei for snRNA-seq data and highlights areas for further research. The effectiveness of a nuclear isolation method for snRNA-seq depends on several factors, including tissue type, processing conditions, and specific research questions. Six samples were sequenced via three methods to observe the methodological variation. Nuclei preparations were repeated multiple times, in some cases dozens of times, to ensure that the sequenced samples were representative of the expected isolation. Although the powering of a single-cell experiment remains a topic of active debate in the field, the proposed approach provides confidence in the results. This study focused on mouse cortical tissue, which offers a specific perspective on the isolation of nuclei. This approach allows for a detailed examination of this particular brain region but may not fully represent the diversity of other brain regions, tissues, or species. Additionally, this study examined the isolation of nuclei under normal physiological conditions. Different pathological states can affect isolation efficacy and quality. Despite these considerations, the findings of this study provide a solid foundation for further research and method refinement in neuroscience and single-cell genomics.

## Supporting information

Supplemental Table 3 - UCell Summary Scoring

Supplemental Table 2 - Microglia Markers

Supplemental Table 1 - Cluster Markers

## ACKNOWLEDGEMENTS

This work was supported by National Institutes of Health Grants R01 AG077829, R01 AG071281, RF1 AG074543, and R21 AG072738 (J.K.); T32AG071444 (to D.J.A.); and the Eli Lilly Stark Neuroscience Fellowship (to L.C.D.). The Kim laboratory was also supported by the Strategic Research Initiative (Indiana University), Indiana University Precision Health Initiative, Indiana University Pervasive Technology Institute (supported in part by Lily Endowment, Inc.), and Shared University Research grants from IBM, Inc., to Indiana University. Single-nucleus RNA libraries were sequenced at the Center for Medical Genomics at Indiana University School of Medicine, which is partially supported by the Indiana University Grand Challenges Precision Health Initiative and the Indiana Genomic Initiative at Indiana University (INGEN); INGEN is supported in part by the Lilly Endowment, Inc. Several figures were created via paid subscriptions to BioRender.com.

## AUTHOR CONTRIBUTIONS

Contributions to this manuscript were made by D.J.A., L.C.D., and J.K. - conceptualization and study design; D.J.A. and L.C.D. - data acquisition and curation; D.J.A., H.N.K., K.H., R.M., and J.H.P. - data analysis; H.N.K. - data interpretation and manuscript draft; and D.J.A., L.C.D., H.N.K., and J.K. – writing, reviewing and editing the manuscript.

## DECLARATION OF INTERESTS

The authors declare no competing interests.

### Declaration of generative AI and AI-assisted technologies in the writing process

This manuscript was developed with the assistance of generative artificial intelligence (AI) tools for writing and editing. Specifically, ChatGPT (4o) (CITE), Gemini (1.5 Flash) (CITE), and Perplexity (CITE) were employed to enhance the clarity and coherence of the text, refine language usage, and annotate code originally drafted by the authors. Generative AI tools were not used to find citations, perform literature reviews, or draft new analyses/interpretations of data. We, the authors, confirm that all scientific ideas, analyses, and conclusions represent our own original work.

## SUPPLEMENTAL INFORMATION

Supplementary Table 1: Excel file containing cluster markers.

Supplementary Table 2: Excel file containing markers for the identified microglial population.

Supplementary Table 3: Excel file containing UCell summary score values for homeostatic astrocytes, microglia, and oligodendrocytes, and disease-associated astrocytes and microglia.

## DATA AVAILABILITY

The snRNA-seq data generated in this study have been deposited in the GEO database (GSE290858).

## CODE AVAILABILITY

The code for the snRNA-seq analysis in this study has been deposited at https://github.com/jungsukimlab/NucleiIsolation.

## METHODS

Key resources table

| REAGENT or RESOURCE | SOURCE | IDENTIFIER |
| --- | --- | --- |
| Chemicals, peptides, and recombinant proteins |  |  |
| EZ nuclei suspension buffer | Sigma | EZ PREP NUC-101 |
| RNAse Inhibitor | Millipore Sigma | 3335399001 |
| Nuclei PURE 2M Sucrose Cushion Solution | Sigma | S9308 |
| Nuclei PURE Sucrose Cushion Buffer | Sigma | S9058 |
| Nuclei Debris Removal Stock Solution | S2 Genomics | #100-253-628 |
| Nuclear Storage Reagent | S2 Genomics | #100-063-623 |
| Critical commercial assays |  |  |
| Chromium Nuclei Isolation Kit | 10X Genomics | PN-1000493 |
| Deposited data |  |  |
| snRNA-seq data | GEO database | GSE290858 |
| Experimental models: Organisms/strains |  |  |
| C57BL/6J | The Jackson Laboratory | 000664 |
| Software and algorithms |  |  |
| CellRanger | 10x Genomics |  |
| SoupX | https://doi.org/10.1093/gigascience/giaa151 |  |
| DoubletFinder | https://doi.org/10.1016/j.cels.2019.03.003 |  |
| Seurat | https://doi.org/10.1038/s41587-023-01767-y |  |
| SCTransform | https://doi.org/10.1186/s13059-019-1874-1 |  |
| scMCA | https://doi.org/10.1007/978-1-4939-9057-3_6 |  |
| PanglaoDB | https://doi.org/10.1093/database/baz046 |  |
| ROGUE | https://doi.org/10.1038/s41467-020-16904-3 |  |
| SAHA | https://doi.org/10.5281/zenodo.14040698 |  |
| UCell | https://doi.org/10.1016/j.csbj.2021.06.043 |  |
| Other |  |  |
| Code | Github | jungsukimlab/NucleiIsolation |

### Experimental model and study participant details

#### Animals

All animal experiments were approved and performed in compliance with the guidelines of the Institutional Animal Care and Use Committee at Indiana University. Male C57BL/6J mice (B6; The Jackson Laboratory, 000664) were maintained under a 12-h/12-h light/dark cycle in a temperature-controlled room with free access to food and water until they reached 6 months of age.

### Method details

#### Tissue collection

At 6 months, B6 mice were anesthetized with Avertin (250 mg/kg, intraperitoneal) and transcardially perfused with cold phosphate-buffered saline (1x PBS). The brains were immediately removed. After the cerebellum and olfactory bulb were removed, the right hemisphere was dissected into cortical and hippocampal regions and immediately frozen in dry ice. The samples were stored at −80°C until further processing.

#### Centrifugation-based nucleus isolation

Frozen anterior cortex tissue was added to a 2 mL tube with 1 mL of ice-cold EZ nuclei suspension buffer (Sigma, EZ PREP NUC-101). The tissue was dissociated via gentle, manual pipetting. The dissociated tissue was transferred into a 15 mL conical tube, and an additional 6 mL of ice-cold EZ nuclei suspension buffer was added. The mixture was incubated at 4°C for 10 minutes while rotating. After rotation, the sample was centrifuged at 500×g for 5 minutes at 4°C. The supernatant was removed, and the pellet was resuspended in 1 mL of ice-cold EZ nuclei suspension buffer with gentle pipetting. An additional 6 mL of EZ nuclei resuspension buffer was added. The mixture was incubated at 4°C for 10 minutes while rotating. After rotation, the sample was centrifuged at 500×g for 5 minutes at 4°C. The supernatant was removed. One milliliter of Nuclei Wash and Resuspension Buffer (NWR Buffer) (1x PBS, 2% bovine serum albumin (BSA), and 2 U/μl RNAse inhibitor (Millipore Sigma, 3335399001)) was added, and the mixture was gently pipetted. An additional 6 mL of NWR Buffer was added. The sample was subsequently centrifuged at 500×g for 5 minutes at 4°C. This wash step with NWR Buffer was repeated four times. After the last wash, the nuclei were resuspended in 1 mL NWR buffer, and then another 6 mL was added. A 70-μm and 30-μm cell strainer was prewetted with LS Column Calibration Buffer (1x PBS, 0.5% BSA). The nuclei were filtered through a 70 μm strainer and then through a 30 μm strainer to remove clumps. The remaining solution was centrifuged at 500×g for 5 minutes at 4°C. The supernatant was removed, and the pellet was resuspended in 1 mL of NWR Buffer. 500 μL of the resuspended nuclei were added to an Eppendorf tube containing 900 μL of sucrose cushion buffer (Nuclei Pure 2 M sucrose cushion solution (Sigma, S9308) and 10% nuclei PURE sucrose cushion buffer (Sigma, S9058)). The mixture was carefully pipetted together. In a separate Eppendorf tube, 500 μL of sucrose cushion buffer was added. To create the sucrose gradient, 1400 μL of the nuclear suspension in sucrose cushion buffer was added carefully to the top of an Eppendorf tube containing 500 μL of sucrose cushion buffer without mixing. The sucrose gradient was centrifuged at 13,000×g for 45 minutes at 4°C. After centrifugation, the supernatant was almost completely removed, leaving approximately 100 μL in the tube. The pellet was resuspended in 1 mL of NWR Buffer with gentle pipette mixing. The solution was filtered through a 40 μm cell strainer to remove any remaining debris.

#### Column-based isolation

Nuclei suspensions were prepared according to the 10x Chromium Nuclei Isolation Kit instructions, with all steps performed on ice[42]. Frozen anterior cortex tissue was added to a prechilled sample dissociation tube (10x Genomics, 2000564). 200 μL of lysis buffer (10x Genomics, 2000558) was added to the tube, and the tissue was dissociated with a pestle. Then, 300 μL of additional lysis buffer was added, and the sample was incubated on ice for 10 minutes. The solution was then added to a Nuclei Isolation Column (10x Genomics, 2000562) in a collection tube (10x Genomics, 2000563). The column was subsequently centrifuged at 13,000×g for 20 seconds at 4°C. The flowthrough was quickly vortexed. The mixture was then centrifuged at 500×g for 3 minutes at 4°C. The supernatant was removed, and the pellet was resuspended in 500 μL of Debris Removal Buffer (10x Genomics, 2000560). The suspension was subsequently centrifuged at 700×g for 10 minutes at 4°C. The supernatant was removed and resuspended in 1 mL of wash and resuspension buffer (1x PBS, 10% BSA, 2.5% RNAse Inhibitor (10x Genomics, 2000565)) and then centrifuged at 500×g for 5 minutes at 4°C. The washing step was repeated once more. The supernatant was removed, and the nuclear suspension was resuspended in 50 μL of wash and resuspension buffer by gentle pipette mixing.

#### Machine-assisted nuclei isolation

Nuclei were isolated via the Singulator 100 system, with all steps performed on ice[43]. Before isolation, nuclei isolation cartridges (S2 Genomics, #100-063-287) were precooled to -25°C, and the simulator was prechilled to ensure optimal processing conditions. For each sample, the frozen anterior cortex was placed into a precooled nucleus isolation cartridge along with 75 μL of RNase inhibitor (Millipore Sigma, 3335399001). The samples were processed via the Standard Nuclei Isolation Protocol. The resulting 3 mL of the nuclear suspension was transferred to a 15 mL conical tube and centrifuged at 500×g for 5 minutes at 4°C to pellet the nuclei. The supernatant was aspirated, and the pellet was resuspended in 3 mL of 20% Nuclei Debris Removal Stock Solution (S2 Genomics, #100-253-628) in Nuclear Storage Reagent (S2 Genomics, #100-063-623). The suspension was centrifuged for 8 minutes at 700×g and 4°C with a decreased brake setting (3 out of 10, Sorvall Legend X1R Centrifuge (Thermo Fisher)). The tubes were carefully removed without disturbing the gradient, and the gradient was aspirated from just below the meniscus via a P1000 pipette to remove the myelin debris, which was visible as a haze or flake at the top. Finally, the nuclei were resuspended in 3 mL of resuspension buffer (2% BSA in Nuclear Storage Reagent).

#### Library preparation

Nuclei concentration and viability were quantified via trypan blue staining and observed with an EVOS XL Core microscope. All the nuclear suspensions were processed with 10x Chromium. Each nuclear suspension was counted and diluted to 1,000 nuclei per μL then loaded into a single-cell chip G and run on the Chromium Controller for GEM generation and barcoding. Sample processing and library preparation were performed according to the manufacturer’s instructions via the Chromium Next GEM Single Cell 3′ v3.1 dual index kit (10x Genomics) and SPRIselect paramagnetic bead-based chemistry (Beckman Coulter Life Sciences). The cDNA and library quality were assessed via a 2100 Bioanalyzer and a high-sensitivity DNA kit (Agilent Technologies). The final library concentration was determined via a QuBit fluorometer and a dsDNA HS assay kit (Thermo Fisher Scientific). Sequencing was performed on a NovaSeq 6000 (v1.5 S2; Illumina) with a 28-10-10-91 read setup to a mean sequencing depth of 437.9 +/-23.9 million reads per sample.

### Quantification and statistical analysis

#### snRNA-seq data analysis

The sequencing data were processed with the Cell Ranger pipeline (v7.0.1, 10x Genomics) and aligned to GRCm 38 (cellranger reference genome: gex-mm10-2020-A).[61]. The filtered feature-cell barcode matrices (including the hashtag count matrix) generated by CellRanger were loaded into SoupX (v1.6.2) in RStudio (v1.4.1717) running R (v4.3.1)[37]. SoupX was used to quantify ambient RNA using default parameters. Doublets were identified with DoubletFinder (v2.0.4)[38]. The data were then loaded into Seurat (v5.0.1)[62]. The data were normalized via SCTransform (v0.4.1)[40]. Nuclei were clustered via the first 35 principal components based on an elbow plot. Cluster marker genes were identified via the *FindAllMarkers* function. Clusters were annotated via scMCA (v0.2.0), which is provided by the Mouse Cell Atlas, and with PanglaoDB[44, 45]. The cell type proportions were quantified using speckle (v0.0.3)[63]. Population homogeneity was estimated with ROGUE (v1.0)[46]. The mitochondria-derived reads were identified by the “mt” prefix, and the ribosomal reads were identified by the “Rp[sl]” prefix. Summary scoring was performed with UCell (v2.2.0)[54].

Figure 1A was created using https://BioRender.com.

